# Liposomal encapsulation of L-arginine and L-citrulline enhances therapeutic effects in a rat model of Preeclampsia and Fetal Growth Restriction

**DOI:** 10.1101/2025.07.21.666044

**Authors:** Caren M. van Kammen, Myrthe J. Brink, Magdalena Minnion, Martin Feelisch, Schiffelers Raymond, A. Titia Lely, Fieke Terstappen

## Abstract

**Background:** L-arginine and L-citrulline improve vascular health in Preeclampsia (PE) and Fetal growth restriction (FGR). However, the short half-life of these amino acids limits their efficacy. This study investigates pharmacokinetics, delivery, and therapeutic outcomes of liposomal encapsulation of L-arginine and L-citrulline in a rat model of PE and FGR.

**Methods:** Firstly, the pharmacokinetics of liposome-encapsulated L-arginine (Encapsulated L-arg) was compared to that of free L-arginine (Free L-arg) in normal pregnant (NP) dams, following single intravenous administration. Secondly, the therapeutic effect on maternal blood pressure and fetal weight were studied in NP rats and the reduced uterine perfusion pressure (RUPP) model for PE and FGR. Treatment groups consisted of Encapsulated L-arginine and L-citrulline (Encapsulated AAs); ratio 1:1, Free L-arginine and L-citrulline (AAs), or PBS, administered intravenously for five consecutive days. Blood and organs were analyzed for amino acid concentrations and fluorescence to assess the biodistribution profile of the liposomes. Nitrite and nitrate were quantified to measure changes in endogenous nitric oxide production in the pharmacokinetic study.

**Results:** Liposomal encapsulation increased the area under the curve of blood arginine concentration-time curves >120-fold. Encapsulated AAs led to a marked increase in plasma and placental tissue concentrations compared to their free forms. Encapsulated AAs reduced maternal blood pressure in RUPP without affecting fetal weight. Enlarged spleens were observed in both Encapsulated AAs groups.

**Conclusions:** The enhanced pharmacokinetics enabled by liposomal encapsulation effectively increased placental delivery of L-arginine and L-citrulline and reduced hypertension. Optimizing composition could enhance efficacy on FGR, making Encapsulated AAs a promising strategy for managing PE and FGR. The etiology of the observed maternal splenomegaly observed warrants further research before this novel approach can be clinically implemented.

## Introduction

Preeclampsia (PE) and fetal growth restriction (FGR) have significant implications for maternal and fetal health. Due to the complex and multifactorial nature of these pregnancy-related complications and the potential risk of any intervention on the developing fetus to date no effective therapeutic approaches for prevention or treatment exist. Reduced nitric oxide (NO) bioavailability plays a key role in maintaining vascular function and placental health. Reduced levels of NO contribute to the pathophysiology of PE and FGR by causing endothelial dysfunction and impaired placental perfusion, compromising nutrient supply. Restoring NO availability represents a promising therapeutic approach to improve maternal and fetal outcomes.

Amino acid supplementation has been extensively studied as a safe option to increase NO availability. Meta-analyses confirmed their favorable effects in preclinical and clinical settings in healthy pregnancies and pregnancies complicated by PE and FGR [1–3]. Many studies have primarily focused on the preventive effects of amino acids such as L-arginine and L-citrulline in PE and FGR through early administration. However, there is growing evidence, particularly with continuous or repeated administration, supporting their therapeutic effectiveness even after PE/FGR onset [4].

L-arginine functions as a substrate for endothelial nitric oxide synthase (eNOS) and is required, together with oxygen, NADPH and tetrahydrobiopterin, to enzymatically produce NO and has been demonstrated to improve blood pressure regulation and support fetal growth in preclinical and clinical studies [2]. L-citrulline, which is produced together with NO from arginine in the same reaction, appears to have comparable effects on FGR [5,6], presumably following recycling to L-arginine, but not on PE, although available clinical studies are limited [7]. L-citrulline may be more effective in bypassing certain regulatory pathways that limit L-arginine availability, particularly in conditions where the endogenous NOS inhibitor asymmetric dimethylarginine (ADMA) is elevated and competes with L-arginine. Combining these two amino acids could provide synergistic effects and offer a more effective strategy to improve maternal, placental, and fetal health [8,9].

Although beneficial therapeutic effects have been demonstrated, maintaining consistent outcomes is challenged by the need for high concentrations and frequent dosing due to rapid clearance. This leads to variable and short-lived effects, making treatment less practical and potentially less effective for patients. Drug delivery systems such as liposomes are known to enhance tissue targeting, pharmacokinetics, and therapeutic efficacy [10,11]. Liposomes are vesicles consisting of one or more phospholipid bilayers enclosing an aqueous core, which can encapsulate hydrophilic compounds such as amino acids, to potentially overcome pharmacokinetic limitations such as the short half-life of promising prenatal therapies, such as L-arginine and L-citrulline.

The current proof-of-concept study aims to explore the potential of liposomal Encapsulated L-arginine and L-citrulline over Free L-arginine and L-citrulline to enhance bioavailability and explore their therapeutic effect in the reduced uterine perfusion pressure (RUPP) rat model.

## 1. Materials and Methods

### 1.1. Encapsulation of the amino acids and quality control

We prepared the liposomes encapsulating L-arginine, alone or in combination with L-citrulline, using the extruder method [12]. The required amount of lipids and fluorescent label DiD 0.1 mol% were dissolved in ethanol and evaporated in a round-bottom flask. The lipid mixture contained Dipalmitoylphosphatidylcholine (DPPC), cholesterol, and poly (ethyleneglycol)-distearoylphosphatidylethanolamine (PEG-DSPE) (molar ratio 1.85 : 1: 0.15). The lipid film was purged with nitrogen to remove residual solvents. Liposomes were formed by rehydration using an aqueous solution containing either L-arginine (0.57 M) or 1:1 of both L-arginine and L-citrulline (each 0.285 M) at a final lipid concentration of 100 mM. To form homogeneous unilamellar liposomes, extrusion under high pressure was performed in a lipex extruder through Anodiscs with 100 nm pore size (Whatman filter, Merck, Darmstadt, Germany). Liposomes were dialyzed overnight against a 1000-fold excess of PBS (Merck) in a 10.000 MWCO membrane tubing (Spectra/Por, Merck). Equal amounts of L-arginine and L-citrulline (0.57 M) were dissolved in sterile water (B. Braun, Oss, the Netherlands) for the Free AAs treatment.

To assess stability, the size and polydispersity index (PDI) were measured with Dynamic Light Scattering (DLS), using a Zetasizer Nano S (Malvern Panalytical, Malvern, UK) for over a month. Liposome samples were diluted 1:50 in H_2_O and measured in three replicates at 25 ^°^ C. Zeta potential was measured by laser Doppler electrophoresis on a Zetasizer Nano Z (Malvern Panalytical). Samples were diluted in 0.1 × DPBS and measured in three replicates at 25°C.

High-Pressure Liquid Chromatography (HPLC) was performed to determine the encapsulation efficiency of the liposome. First, with Bligh and Dyer extraction separation the lipid compounds from the liposomes were separated from the samples to prevent blocking of the HPLC [13]. The aqueous phase was used for HPLC analysis. The encapsulation efficacy was calculated by Encapsulation Efficiency (%) as the amount of AA in liposome divided by the total amount of AA × 100%.

We evaluate drug leakage *in vitro* with a dialysis assay by placing a 10,000 MWCO Slide-A-Lyzer cassette (Thermo Fisher Scientific, USA) containing liposome sample in 80 ml PBS at 37 °C on an orbital shaker. Samples were collected from PBS every 24h for five days and measured with HPLC to measure the amount of AAs leaking out of the liposome.

### 1.2. Ethical considerations

This study was conducted following institutional guidelines for the care and use of laboratory animals of Utrecht University and the University Medical Center Utrecht, and all animal procedures were approved by the local Animal Welfare Body (AWB; Utrecht, the Netherlands) under an Ethical license provided by the national competent authority (Central Committee on Animal Experiments, CCD, The Netherlands), securing full compliance with the European Directive 2010/63/EU for the use of animals for scientific purposes. Prior to the *in vivo* study, a power calculation was performed in G-power. Ten animals per group were determined to reach the desired effect size of 1.46, with alpha of 0.0167, and a power of 0.8 of a clinically relevant difference of Mean Arterial blood Pressure (MAP) (primary outcome) of 5 mmHg. All efforts were made to design and report the animal experiments according to the ARRIVE guidelines (**Table S1**).

### 1.3. Animals

Twelve time-pregnant virgin Sprague Dawley rats between twelve and thirteen weeks old were purchased by Envigo (Envigo RMS B.V, Horst, the Netherlands) for a pharmacokinetic study, and a total of 66 were ordered in three cohorts to study the therapeutic effect. Dams were purchased at gestational day (GD) 4, 5, 6, or 7 and housed in groups in conventional cages for acclimatization to local vivarium conditions. Animals were kept under a regular 12h light/dark cycle at a room temperature of 22 °C and relative humidity of 50%. At GD13, dams were housed individually in cages to measure daily water and food intake. All animals had free access to food (Ssniff R/M maintenance, Germany), and water.

To study the pharmacokinetic differences in normal pregnant (NP) dams between those randomly assigned to Encapsulated L-arginine (Encapsulated L-arg), Free L-arginine (Free Arg), or vehicle (phosphate-buffered saline, PBS) using a computer-based random order generator [11]. To study the therapeutic effect, dams were randomized into reduced uterine perfusion pressure (RUPP) or normal pregnant (NP) group and further randomly assigned to one of three treatment groups: Encapsulated AAs, Free AAs, or PBS as a control.

In both studies, caregivers were blinded; for researchers, blinding was not feasible for the group assignment and intervention, as group allocation was visibly apparent based on factors such as body weight and the presence of surgical wounds. Additionally, in the intervention group, the Encapsulated AAs showed a bluish coloration, while the Free AAs, or PBS, appeared white, further preventing effective blinding. Blinding on outcome measures was addressed by ensuring that researchers were unaware of the treatment allocation during data analyses.

### 1.4. Pharmacokinetics of liposome-encapsulated L-arginine

The NP dams were administered intravenously with a single dose of either Encapsulated L-arg, 15mg/kg, Free L-arg, 15mg/kg, or PBS at GD17. To determine the amino acid concentration and NO in plasma and organs, blood samples were collected in an EDTA tube via the Saphenous vein puncture, with a maximum of 6 timepoints per animal to adhere to animal welfare guidelines. Sampling time points were selected to capture both expected rapid (Free L-arg) and slower (Encapsulated L-arg) clearance differences between groups, with overlapping time points at t= 0 min before injection, after injection 1 h, 24 h, and 48 h. Extra timepoints for PBS: 30 min, 4h; Free L-arg: 5 and 30 min; Encapsulated L-arg: 4, 8, and 30 h. The last sample of all dams was collected via cardiac puncture right before euthanasia at 48 hours. Experimental concentration data were fitted to the exponential decay model using GraphPad Prism (Boston, MA, USA). Before model fitting, endogenous L-arginine concentrations from the PBS control group were subtracted from the Free L-Arg and Encapsulated L-arg data to account for baseline levels.

### 1.5. Therapeutic effect of liposome-encapsulated AAs

The dams were intravenously administered daily from GD15 to GD19 with Encapsulated AAs 10 mg/kg, Free AAs 10 mg/kg, or PBS. Blood was collected in an EDTA tube from the dams by puncturing the lateral saphenous vein every 24 hours and immediately centrifuged for 3 minutes at 16000 x g to separate the erythrocytes from plasma. This was stored on dry ice and transferred to −80 °C storage.

### 1.6. Induction of PE/FGR model

The Reduced Uterine Perfusion Pressure (RUPP) surgery was performed at gestational day fourteen [14]. In short, under isoflurane anesthesia (3% induction, 2–2.5% maintenance), a midline abdominal incision exposed the uterine horns and uteroplacental vessels. For RUPP surgery, a silver clip (AG000450; 1.5 cm × 15 mm, 0.203 mm inner diameter) was placed around the abdominal aorta above the iliac bifurcation. Two smaller clips (1 cm × 15 mm, 0.100 mm inner diameter) were placed on the right and left ovarian arteries to block compensatory blood flow. Animals received subcutaneous carprofen (5 mg/kg) 30 minutes before and 24 hours after surgery for analgesia.

### 1.7. Carotid artery cannulation

Cannulation of the carotid artery was conducted to measure mean arterial blood pressure (MAP) at GD18. A small ventral midline neck incision from the lower part of the mandible to the sternum was made. Next, the left common carotid artery was isolated. A small incision was made, followed by insertion of a catheter tip reaching the aortic arch. The catheter was subcutaneously transferred and exteriorized between the shoulder blades. The catheter was flushed with heparinized saline (300 mg/ml, Heparin LEO 5.000 IU, LEO Pharma B.V, Amsterdam, the Netherlands) to reduce the risk of blood clotting around the catheter. The incision in the neck was closed using surgical tissue adhesive (3 M™ Vetbond Tissue Adhesive 1469, 3 M, Saint Paul, MN, USA). The animals received a subcutaneous injection of carprofen (Zoetis Inc., Parsippany, NJ, USA) (5 mg/kg body weight) as an analgesic 30 minutes before surgery and 24 hours post-operatively.

### 1.8. Blood pressure measurements and analyses

To reduce stress during the conscious blood pressure measurements, the dams were trained beforehand in the fixation tubes on three separate occasions to ensure conscious and stable measurements. On GD19, the mean arterial pressure (MAP) and heart rate were recorded in a computer-generated random order. Dams were secured in fixation tubes, and the carotid artery catheters were connected to the pressure transducer. A three-way valve was placed on the line to flush the cannula with heparinized saline (300 mg/ml, Heparin LEO 5.000 IU, LEO Pharma B.V) before measurements. The signal was amplified (PowerLab 8/35, ADInstruments, Dunedin, New Zealand), digitized, and recorded with LabChart Pro v8. 1. 19 (ADInstruments). A habituation period of 30 minutes was implemented to acclimate the dams before the start of the measurement. Heart rate values of ≤300 bpm and ≥600 bpm were considered artifacts and excluded from the blood pressure analysis.

### 1.9. Euthanasia and organ collection

In both experiments, the dams were euthanized under isoflurane anesthesia by performing a heart puncture, resulting in total bleeding at GD19. The dams received carprofen (5 mg/kg body weight, s.c.) as an analgesic at least 30 minutes before euthanasia. Subsequently, animals were perfused using ice-cold phosphate-buffered saline (PBS). Urine samples were obtained by bladder puncture. The maternal liver, maternal kidneys, maternal spleen, placentas, fetuses, and fetal livers were removed and weighed. The organs were immediately snap-frozen in liquid nitrogen and stored in a −80 °C freezer. For the LC-MS/MS analysis of the amino acids of interest and the NO oxidation product analysis, organs were crushed into a fine powder and stored again in a −80 °C freezer before further processing.

### 1.10. LC-MS/MS analysis of amino acids in tissue and blood plasma

L-arginine and L-citrulline concentrations, together with other amino acids (**Table S2**), play an essential role in the endogenous formation of NO, and were measured using Hydrophilic interaction chromatography (HILIC) combined with tandem mass spectrometry (MS/MS) (HILIC-MS) according to Prinsen [15]. In short, the organ samples were diluted 1:10 in 100% methanol and further homogenized using a bullet blender tissue homogenizer for 10 min at 4 °C. After centrifuging at 16,200 x g for 5 min, the supernatant was stored at −80 °C. For plasma and tissue analyses, 20 μL plasma (or tissue extract) were mixed with 20 μL of internal standard (IS) solution and were run over a Acquity UPLC BEH Amide column (2.1×100 mm, 1.7 μm particle size) including a Van Guard™ UPLC BEH Amide pre-column (2.1×5 mm, 1.7 μm particle size), using a Waters ACQUITY UPLC system (Waters, Milford, USA). Plasma samples with high AAs concentrations were diluted until the concentrations of the AAs fell within the range of the calibration curve.

The column was maintained at 35 °C with a 0.4 mL/min flow rate. A gradient was applied using solvent A (10 mM ammonium formate in 85% acetonitrile with 0.15% formic acid) and solvent B (10 mM ammonium formate in Milli-Q water with 0.15% formic acid, pH 3.0). The first 6 minutes were 100% solvent A, followed by a 0.1 min gradient to 94.1% A / 5.9% B. Over the next 3.9 min, solvent A decreased to 82.4% and B increased to 17.6%, then adjusted to 70.6% A / 29.4% B for the final 2 min. The column was re-equilibrated for 6 min, making a total run time of 18 min. The column was coupled to a triple-quadrupole mass spectrometer (Waters Xevo TQ) in ESI-positive mode with a capillary voltage of 1.00 kV, desolvation temperature of 550 °C, source temperature of 150 °C, cone gas flow of 50 L/h, and desolvation gas flow of 1000 L/h.

### 1.11. HPLC analysis of nitrite and nitrate as markers of NO production in blood, tissue, and urine

To assess whether L-arg or the L-arg/cit mixture affected endogenous NO formation, its major oxidation products nitrite and nitrate were analyzed in representative organs, blood plasma and urine. After dilution of pulverized organ homogenates with ice-cold PBS (1:5, weight/volume), aqueous tissue extracts were processed further using a Potter-Elvehjem homogenizer (350 rpm/minute with 6.5 up/down strokes three times while immersed in crushed ice. Homogenates were stored in a −80 °C freezer until further analysis. Tissue homogenates were deproteinized with methanol (volume was matched to the added volume of PBS used to homogenize the tissue samples). After vortexing for 1 min and high-speed centrifugation (21,100 × g for 20 min at 4°C), the supernatant was injected onto the HPLC system.. Aliquots of the thawed Red blood cell pellets were weighed into fresh Eppendorf tubes and the appropriate volume of lysis solution (5mM EDTA and 10mM N-ethylmaleimide in MilliQ water) was added (ratio 1: 4, weight: volume). The samples were sonicated with an ultrasonic probe for 30 seconds and then centrifuged to separate the supernatant from the ruptured cell membrane. Protein was removed by methanol precipitation (1:1, v/v) as described for tissue homogenates. An aliquot of dry rodent chow was weighed and suspended in PBS (1:5, weight/volume), vortexed and kept frozen at −80 °C until analysis at which stage protein was removed by methanol precipitation (1:1, v/v), as for all other specimen. All samples were analyzed for nitrite and nitrate by high-pressure liquid chromatography using a dedicated analysis system (ENO-30 analyzer with AS-700 INSIGHT autosampler and Clarity Envision software; Amuza, Inc, San Diego, CA/Eicom, Japan).

### 1.12. Statistics

All statistical analyses were performed using GraphPad Prism v10.0.2 for Windows (GraphPad Software, Boston, MA, USA). The experimental unit in this study is the dam, as the inducement and treatment are applied at the maternal level, and litter outcomes are considered as dependent observations. In the pharmacokinetic study, a one-way ANOVA was conducted to analyze differences among the three treatments within the NP group, as the samples were independent and continuous. For the therapeutic effect study, we examined two distinct groups, RUPP and NP, each subjected to three different treatments. Differences between the two groups and three treatments were investigated with a two-way ANOVA with a post-hoc multiple comparisons test if applicable. Data are predominantly reported as mean ± standard deviation (SD); only L-arginine concentrations in the PBS group are presented in text as median (minimum–maximum) values. The results were considered significantly different with a two-sided p-value < 0.05.

### 1.13. Exclusion criteria and dropouts

Animals were excluded from participation in the experiment if they were not pregnant at delivery or after surgery or if they reached the humane endpoint before the end of the experiment. We excluded dams from the study when immediate humane endpoints were reached in cases of severe distress, such as lethargy, abnormal rattling breathing, complications during or after surgeries (RUPP or carotid artery), such as fully open surgical wound, and paralysis.

To reduce heterogeneity, we standardize exclusion criteria. Dams were excluded from the study for all outcomes with a cut-off value of litter size <4 or >16, as very small or large litter sizes increase variability in fetal weight outcome [16]. Exclusions and final sample size of the different phenotypical outcome parameters are presented in **Table S3**.

## 2. Results

### 2.1. Characterization of liposomes

The average particle size measured was 0.14 μm with a polydispersity index of 0.064, which remained stable over at least one month. This shows a high degree of physical stability and a small distribution width, indicating that the liposome population was monodisperse, confirming their structural integrity. Encapsulation efficacy was 12%. This is expected for water-soluble compounds to be around 10% at this liposome concentration. The surface charge was −6.21 mV, which is slightly anionic and near neutral. Dialysis at 37 °C under shaking conditions showed limited leakage of a maximum of 5% of the encapsulated amount over 96 hours.

### 2.2. Liposome-encapsulation of L-arginine Improves pharmacokinetics in healthy pregnant animals without altering steady-state concentrations of NO

In healthy pregnant rats, Encapsulated L-arg or Free L-arg were injected intravenously at a dose of 10 mg/kg, and the pharmacokinetics and tissue concentrations were compared to PBS-injected controls. A dramatic improvement of the pharmacokinetic profile of L-arginine was observed upon liposomal encapsulation (**Figure 1A**). The area under the concentration-time curve was markedly increased for the Encapsulated L-arg group (3074 ± 83 µg/ml^*^ min) compared to the Free AA group (25 ± 1 µg/ml^*^ min). The Encapsulated L-arg group reached a higher maximum concentration (155 ± 19 µM) compared to the Free AA group (34 ± 2 µM). Also, a significantly lower clearance was noted in the Encapsulated L-arg group (0.0016 ± 0.0005 ml/min) compared to the Free AA (0.0681 ± 0.001 ml/min) group. Thereby, the half-life of L-arginine was significantly enhanced in the Encapsulated L-arg (440 ± 120 min) compared to the Free AA group (10 ± 2 min). Before injection, circulating endogenous L-arginine in all three groups was comparable (185 ± 3 µM). Concentrations in the PBS group remained stable over all time points with a median of 168.50 µM, ranging from 103 to 198 µM. Unexpectedly, no significant differences in circulating nitrite and nitrate concentrations were observed between groups (**Figure 1B** and **Table S4**). Likewise, acute increases in L-arginine availability did not alter fetal, placental, and maternal tissue levels of these NO products (**Fig. S1**). Urinary nitrate levels did not differ significantly between groups at 48 hours post-treatment (**Figure 1C**).

**Figure 1.**
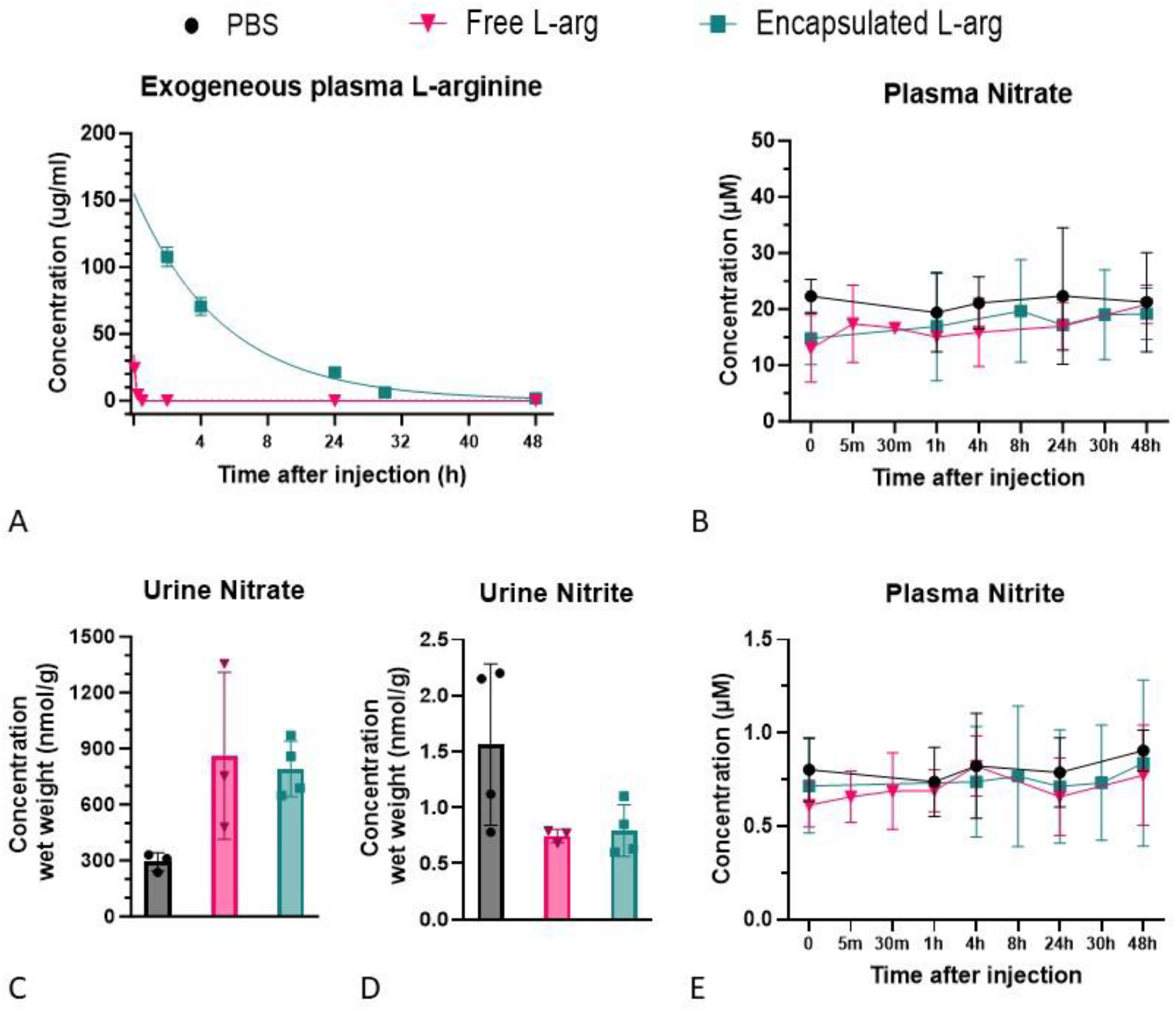
Pharmacokinetics of L-arginine in normal pregnant rats. **A)** Improved pharmacokinetic profile of exogenous plasma L-arginine of Encapsulated L-arginine group in logarithmic scale with estimated C(0) **B)** No differences in plasma concentrations of nitrate **C)** Urinary excretion of nitrate and **D)** nitrite **E)** and plasma concentrations nitrite in animals treated with either PBS, Free L-arg or Encapsulated L-arg (n=4 per group). Except for Urinary nitrate PBS N=3, one animal no data point. Data are depicted as mean ± SD. PBS, Phosphate-Buffered Saline; Free L-arg, Free L-arginine; Encapsulated L-arg, Encapsulated L-arginine.

### 2.3. Enhanced uptake of liposome-encapsulated AAs RUPP model

Based on the results of Encapsulated L-arg, we studied the biodistribution of Encapsulated L-citrulline and L-arginine (Encapsulated AAs) compared to their free forms (Free AAs) in the RUPP rat model of PE/FGR. The Encapsulated AAs group showed higher concentrations of both citrulline and L-arginine relative to free form in blood (**Figure 2A**) and in placental tissue (**Figure 2B**), but not in the fetal liver (**Figure 2C**). Food and water intakes were comparable between groups, indicating a consistent supply of experimental animals with protein (giving rise to arginine and citrulline) (**Figure S1**) as well as intake of nitrite and nitrate from the rodent chow (measured chow nitrate 928 Umol/g and nitrite 7.09 Umol/g. The treatment did not affect other NO-cycle amino acid levels in plasma or tissue (data not shown). Based on DiD fluorescence, a small portion of the injected liposome dose accumulated in the placenta, and no accumulation was observed in the fetal liver (**Figure 2D**), with the highest uptake in the liver (36 ± 10 % per organ) and spleen (19 ± 8 % per organ) (**Figure S2**).

**Figure 2.**
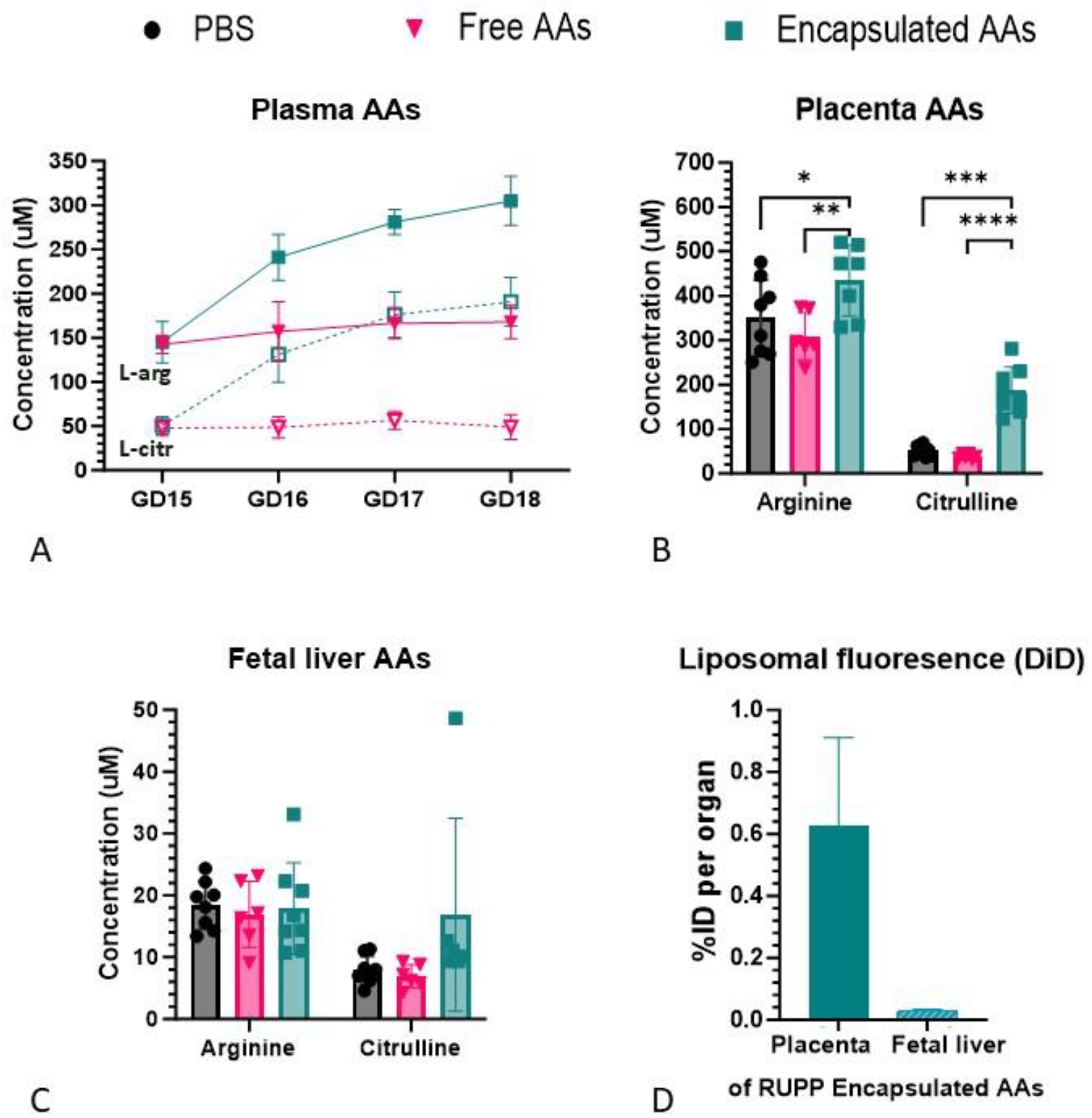
Concentration profiles of L-arginine and L-citrulline in plasma and tissues of RUPP rats. **A)** L-arginine (closed square and triangle) and L-citrulline (open square and triangle) in plasma between Free AAs (magenta, n=7), or Encapsulated AAs (Green, n=7) from GD15-18 **B)** L-arginine and L-citrulline concentration in the placenta at GD19, PBS (n=8), Free AAs (n=7), or Encapsulated AAs (n=7). **C)** L-arginine and L-citrulline concentration in the fetal liver at GD19, PBS (n=8), Free AAs (n=7), or Encapsulated AAs (n=7). **D)** The fluorescent label DiD uptake of liposome in placenta and fetal liver at GD19 of Encapsulated AAs (n=7). Data are depicted as mean ± SD. P value = ^*^ p<0.05,^**^P<0.01,^***^indicates P<0.001, ^****^ P<0.0005. RUPP, Reduced Uterine Perfusion Pressure; PBS, Phosphate-Buffered Saline; Free AAs, Free L-arginine + L-citrulline; Encapsulated AAs encapsulated L-arginine + L-citrulline; L-arg, L-arginine; L-cit, L-citrulline; %ID, percentage injected Dose.

### 2.4. Therapeutic effect of liposome-encapsulated AAs on hypertension and fetal growth

The PE and FGR phenotypes were assessed to evaluate whether the increased pharmacokinetics and uptake of Encapsulated AAs enhance their therapeutic effect on maternal blood pressure and fetal weight compared to Free AAs treatment.

The Encapsulated AAs treatment decreased MAP with a mean of −17 mmHg, whereas Free AAs did not lower MAP compared to RUPP-PBS (**Figure 3A**). No therapeutic effect was observed on fetal weight in RUPP of either treatment group. (**Figure 3B**). Only in the NP group Encapsulated AAs treatment improved fetal weight compared to Free AAs. Mirroring the fetal weight outcome, only in the NP group Encapsulated AAs increased placental weight compared to Free AAs (**Figure 3C**). The reabsorption rate of litter from the dam increased in RUPP-PBS group compared to NP-PBS (**Figure S3A**). There was no improvement in reabsorption rate in both treatment groups compared to RUPP-PBS. Mean litter size (implantation sides) were not significantly different between groups (**Figure S3B**). Encapsulated AAs therapy increased maternal spleen weight in both NP and RUPP (**Figure 3D**).

**Figure 3.**
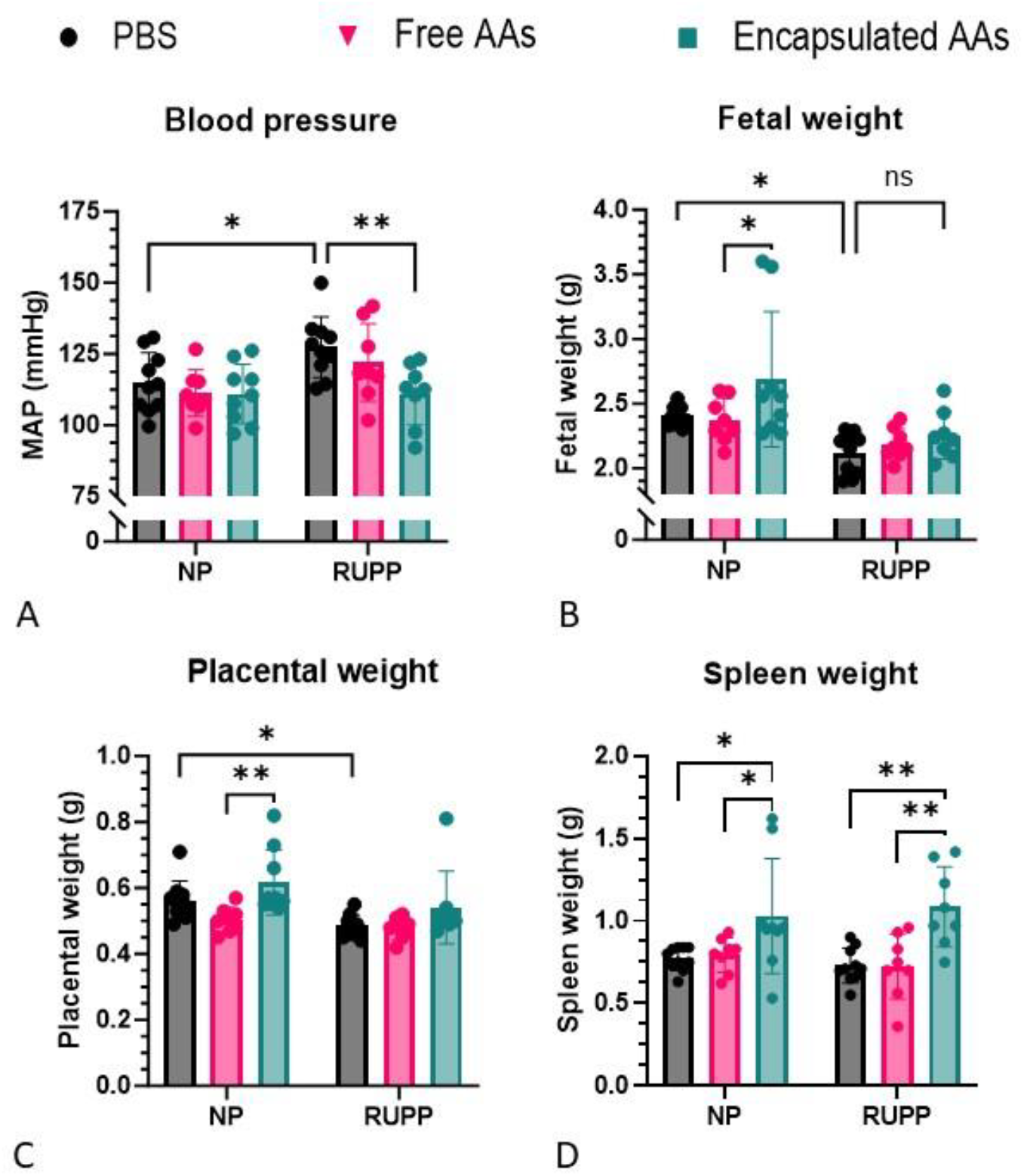
Phenotypical treatment outcomes of Free AAs and Encapsulated AAs at GD19. **A)** Decreased maternal blood pressure of treatment with Encapsulated AAs in RUPP dams **B)** No fetal weight improvements in treatment groups of Free AAs and Encapsulated AAs in RUPP **C)** No placental weight improvements in treatment groups of Free AAs and Encapsulated AAs in RUPP **D)** Enlarged spleen weight in Encapsulated AAs groups in NP and RUPP. Treatment PBS as control (NP n=10 RUPP n=10), Free AAs (NP n=8 RUPP n=8), Encapsulated AAs (NP n=9 RUPP n=8). Data are depicted as mean ± SD. P value = ^*^ p<0.05, ^**^P<0.01,^***^ indicates P<0.001, ^****^ P<0.0005. NP, normal pregnant; RUPP, Reduced Uterine Perfusion Pressure; MAP, mean arterial pressure; PBS, Phosphate-Buffered Saline; Free AAs, Free L-arginine + L-citrulline; Encapsulated AAs, Encapsulated L-arginine + L-citrulline; GD, Gestaional Day

## 3. Discussion

The liposomal encapsulation of L-arginine, alone and in combination with L-citrulline, resulted in an enhanced pharmacokinetic profile and delivery of these amino acids compared to their administration in free form, contributing to the mitigation of high blood pressure in the RUPP model of placental insufficiency-induced PE and FGR. Only NP dams receiving Encapsulated AAs showed increased fetal and placental weights.

### 3.1. Enhanced pharmacokinetics increased therapeutic impact on only blood pressure and optimized delivery to placenta

The prolonged half-life and increased plasma concentration of L-arginine by liposomal encapsulation are consistent with findings from studies on liposomal encapsulation of other water-soluble therapeutics in non-pregnant animals [17–21].

### Impact on maternal outcome

Enhancing systemic L-arginine and L-citrulline availability by liposome-encapsulation effectively improved mean arterial pressure (MAP) in RUPP compared to normal pregnancy in our study, which aligns with preclinical and clinical findings where L-arginine administration alone reduced blood pressure [22–24]. While L-citrulline has been shown to reduce blood pressure in in vivo models of PE and FGR (26, 27), clinical studies in women with hypertensive disorders, including preeclampsia, have reported inconsistent effects [25,26]. In this context, we were unable to distinguish the specific contributions of L-arginine and L-citrulline to MAP changes in our study.

Although in our study the Encapsulated AAs effectively improved MAP, whereas Free AA did not, the BP-lowering effect is not significantly stronger than the Free AAs (p = 0.11). While encapsulation offers the advantage of sustained amino acid delivery [27]. The encapsulation process slows release kinetics, which could reduce immediate bioactivity and delay the acute therapeutic effects in our study. Additionally, the relatively short treatment duration and the model used may not have fully captured the potential benefits of prolonged Encapsulated AAs delivery. Supporting this, another study reports greater effects with Free L-arginine given over a longer period by a combined preventive and therapeutic treatment approach [28], suggesting that extended treatment windows might be beneficial in maximizing the effect size of treatment with Encapsulated AAs. Addressing this requires extending these investigations to other models, such as the Dahl Salt-Sensitive rat with pre-existing hypertension and proteinuria, which involves superimposed preeclampsia during pregnancy [28].

### Impact on fetal and placental outcome

Encapsulated AAs enhanced fetal and placental growth in NP animals, but did not significantly improve outcomes in the RUPP model. This is in line with a study of oral supplementation of Free L-arginine in the same model [29]. In contrast, in other models of FGR, the majority showed positive effects of L-arginine or L-citrulline oral supplementation on fetal and placental weight [5,6,28,30–32]. Only a small fraction of our liposomes crossed the placenta, suggesting this mode of delivery is safe for the fetus. However, effective fetal growth relies more on the delivery of L-citrulline as it acts as a stable precursor that raises fetal L-arginine levels, and is crucial for fetal muscle development. The minimal placental transfer of the exogenous L-citrulline may explain why fetal growth [5,6,28,30].

The Encapsulated AAs significantly increased delivery of L-citrulline in the placenta, suggesting a potential mechanism for improving fetal health through enhanced placental transport of nutrients. Targeting ligands to optimize placental delivery could enhance amino acid exchange at the maternal-fetal interface, improved by local efficacy at the placenta.

### 3.2. Therapeutic safety with caution for immune-altered conditions in PE and FGR

Liposomes are widely used in drug delivery due to their biocompatibility and ability to modify the pharmacokinetics of the cargo. Many different liposomal drugs and vaccines are FDA-approved and generally recognized as safe (GRAS); however, in the present study, we found enlarged spleens after repeated administration of our PEGylated liposomes in both normal pregnancy and pregnancies complicated with PE and FGR. Pregnancy [33,34], particularly when complicated by PE and FGR [35], are known to be associated with modulation of immune function. Repeated exposure to liposomes during pregnancy may further enhance the immune modulatory response. Our liposomes were likely taken up by splenic macrophages, contributing to immune activation and increased workload on phagocytic cells [36]. Although the negative charge (−6 mV) of PEGylated liposomes reduces direct platelet binding compared to cationic liposomes, they could still trigger immune mechanisms that sequester platelets in the spleen indirectly [37,38]. Whereas potential impacts on spleen function and immune activation could negatively affect maternal and fetal health, no adverse effects were observed macroscopically or in the form of alterations in maternal and fetal weight. This suggests that the treatment does not compromise overall maternal or fetal development. However, a thorough investigation into the potential modulation of maternal and fetal immune responses remains warranted.

### 3.3. How may enhanced L-arginine and L-citrulline availability mediate its beneficial effects?

The observed beneficial effects of combined L-arginine and L-citrulline administration on blood pressure in the RUPP model supports the potential of this form of amino acid administration to improve NO bioavailability. If an elevation of systemic L-arginine availability would indeed translate into an improvement in endothelial function by increasing vascular endothelial NO production, one would expect this to be associated with a selective increase in circulating nitrite concentrations in blood plasma. [39,40]. However, this was not observed in our study. Neither did the concentrations of nitrite and/or nitrate change markedly in maternal or fetal organs following acute administration of Free or Encapsulated L-arg in healthy, pregnant rats. This suggests that endogenous NO production at the tissue and whole-body levels may not significantly change in response to acute alterations in circulating levels of its substrate, L-arginine. Moreover, our findings did not detect increased nitrate excretion with both Free and Encapsulated forms. This may be caused by suboptimal timing of nitrate and nitrite measurements, with potentially stronger responses detectable at earlier time points. It remains to be determined whether circulating, urine, and/or tissue concentrations of NO metabolites change when L-arginine and L-citrulline levels are robustly elevated over a prolonged period of time starting earlier.

### 3.4. Strengths and limitations

This is the first proof-of-concept study that improving the pharmacokinetics of promising pharmacological principles with short half-lives can translate into a difference in therapeutic efficacy, as shown here for Encapsulated L-arg treatment compared to Free L-arg and control during pregnancy. Moreover particular strength of our investigation is that we conducted *in vivo* experiments on the phenotypical effects of combined encapsulation of L-arginine and L-citrulline in a robust placental insufficiency-induced PE and FGR model, while evaluating multiple perinatal outcome parameters such as fetal weight, placental weight, and maternal blood pressure in the RUPP model.

However, our studies are not without limitations. The absence of a dose-response study and the role of citrulline-to-arginine ratio and optimal dosing of the AAs remain unclear. L-arginine and L-citrulline mainly impact one pathway of the complex multifactorial PE/FGR pathophysiology, which could mainly improve the associated vascular alterations. Furthermore, the short therapeutic window in the RUPP model restricts the assessment of longer-term treatment effects. Finally, it remains uncertain at this stage whether or not the beneficial effects of enhancing L-arginine availability is mediated via an improvement in endothelial NO production and/or other metabolic effects.

### 3.5. Conclusion and perspectives

The encapsulation of L-arginine and L-citrulline show promising effects on maternal blood pressure as a result of enhanced pharmacokinetics and increased delivery of these aminoacids to the placenta. Collectively, the improved maternal health conditions may prolong pregnancy duration, which in turn contributes to the reduction of potential health risks for the baby associated with premature birth.

While liposomes are designed to address transport issues, they may not fully resolve the underlying problem, as the impaired placental transport of amino acids in FGR still limits the potential for proper fetal development. Thus, a more comprehensive solution is needed to enhance amino acid transfer and improve fetal growth and development and thus pregnancy outcomes.

Future research should focus on investigating the ratio of AAs and integrating complementary pathways, which together could enhance the impact on FGR phenotype. Furthermore, optimizing placenta-targeting by receptor-specific nanobodies allows for lower doses and reduces systemic side effects and immune reactions by directing drug delivery specifically to the placenta instead of off-target organs like the spleen. This strategy may contribute to placental health due to increased efficacy and consequently improve fetal growth. Moreover, the complex relationship among high-risk pregnancies, splenomegaly, and innovative drug delivery technologies poses a challenge that warrants more research as a crucial step towards clinical implementation.

### Novelty and Relevance

#### What Is New?

- Improved pharmacokinetics of L-arginine by liposomal Encapsulation versus Free administration during pregnancy.
- First study on therapeutic efficacy of Encapsulated L-arginine and L-citrulline in placental insufficiency-induced preeclampsia and fetal growth restriction rat model.

#### What Is Relevant?

- Improved maternal blood pressure regulation by elevated efficacy, without affecting placental or fetal outcome.
- In hypertensive disorder of pregnancy, the altered immune modulation and organ function may heighten risks from liposome interactions.
- Highlights the need for optimizing placenta-targeted drug delivery for better therapeutic outcomes.

## Acknowledgments

The successful completion of this research project would not have been possible without the support and contributions of many individuals and organizations. We would like to express our gratitude to the Metabolic Diagnostic Department at UMCU, especially Marjolein Bosma, for their expert assistance with amino acid concentration measurements.

## Supplement material

Tables S1-S4

Figures S1-S3

